# Single-molecule acceptor rise time (smART) FRET for nanoscale distance sensitivity

**DOI:** 10.1101/2023.03.15.532809

**Authors:** Jiajia Guo, Xuyan Chen, Premashis Manna, Xingcheng Lin, Madelyn N. Scott, Wei Jia Chen, Mikaila Hoffman, Bin Zhang, Gabriela S. Schlau-Cohen

**Affiliations:** Department of Chemistry, Massachusetts Institute of Technology, MA, USA; Institute of Biomedical and Health Engineering, Shenzhen Institute of Advanced Technology, Chinese Academy of Sciences, Shenzhen, China; Department of Chemical and Biological Physics, Weizmann Institute of Science, Rehovot, Israel; Department of physics, University of Illinois Urbana Champaign, IL, USA

## Abstract

The structure, dynamics, and binding of individual biomolecules have been extensively investigated using single-molecule Förster resonance energy transfer (smFRET) as a ‘spectroscopic ruler.’ The FRET efficiency between a fluorophore pair is used to measure distances in the several nanometer range. Existing approaches to detect closer distances come at the expense of sensitivity to longer distances. Here, we introduce single-molecule acceptor rise-time (smART) FRET that spans closer and longer distances. The acceptor rise time encodes the FRET rate, which scales polynomially with distance and thus has a steep dependence that expands the working range by 50%. High precision and accuracy is achieved through the spectroscopic separation between the rise time and the photophysical fluctuations that obfuscate other FRET readouts. Using the nanoscale sensitivity, we resolved the architectures of DNA bound to the single-stranded binding protein from *E. coli*, demonstrating the ability of smART FRET to elucidate the complex behaviors of biomolecules.

## Introduction

Visualizing the conformations, interactions, and dynamics of biomolecules is a fundamental and critical challenge in understanding biological processes^1,2^. While advances in high-resolution structural techniques have revealed detailed pictures of biomolecular conformations, the associated dynamics, particularly under physiological conditions, have been more challenging to access^3,4^. Single-molecule Förster Resonance Energy Transfer (smFRET) is a powerful and non-invasive approach to probe dynamics^5,6,7^. In smFRET, intra- and intermolecular distances for individual biomolecules are measured through the efficiency of energy transfer between two conjugated fluorophores^8,9^. Typically, the efficiency is calculated from the relative fluorescence intensities of the donor and acceptor fluorophores and then converted into distance^4, 10^. However, the working range of intensity-based measurements is fundamentally limited to interfluorophore distances for which the timescale of energy transfer is similar to the fluorescence lifetime of the donor fluorophore. Outside of this limited range, the intensity approaches zero for the donor or acceptor fluorophore at shorter or longer distances, respectively. At low intensities, small efficiency changes are challenging to detect at the single-molecule level due to the low signal-to-noise and signal-to-background ratio of the data^11^. Methods using fluorescence quenching or charge transfer have been developed to extend the working range down to short distances, but these advances come at the expense of longer distances and require a new toolkit of labels^12,13,14^. These challenges have collectively limited the range of intensity-based smFRET measurements. Many biological systems rely on nanoscale interactions, yet the current range of smFRET cannot resolve these key interactions and how they are modulated to drive biological function.

As an alternative to intensity-based measurements, the timescale of energy transfer can be directly measured and converted to efficiency. The timescale has a steep dependence over a wide range as it scales polynomially with distance, *i*.*e*., 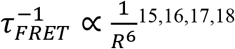. Over the nanoscale range relevant for smFRET, the timescale changes by seven orders of magnitude. To date, timescale-based methods have used only the fluorescence lifetime of the donor ^19, 20^. However, at low efficiencies, fluctuations in the donor photophysics obscure small efficiency changes and, at high efficiencies, the donor signal is too low to accurately fit the lifetime. Furthermore, background emission is strongest in the donor channel and can obfuscate the donor decay. Collectively, these challenges lead to an operating range with similar constraints as intensity-based methods^20^.

The timescale of energy transfer is also encoded in the rise of the acceptor fluorescence, a readout spectroscopically separated from photophysical fluctuations in intensity or lifetime and from background emission. Single-molecule measurements of the rise were not possible until the recent development of fast single-photon detectors^21^. Using this technology, we introduce single-molecule acceptor rise time (smART) FRET, a method that extracts the efficiency from the acceptor rise. smART FRET outperforms other FRET approaches with a 50% expansion of the distance range and significant improvements to the accuracy. In particular, smART FRET maintains accuracy and precision in the high FRET regime that captures short distances. Finally, smART FRET utilizes the same toolkit of labels as traditional FRET methods, and thus is applicable to the same wide range of proteins, peptides, and nucleic acids of biophysical interest. Here, we demonstrate the accuracy and versatility of smART FRET using a DNA ruler control and the single-stranded binding protein from *E. coli*.

## Results

### smART FRET on DNA rulers

The performance of smART FRET was quantified and compared to traditional FRET methods by measuring DNA duplexes with Atto550 and Atto647N attached as a donor and acceptor pair separated by a distance range of 3.77 to 8.66 nm, 1.5 times wider than other smFRET benchmarking studies^10^. Single-molecule fluorescence was recorded for duplexes immobilized on a coverslip using a confocal microscope with time-correlated single photon counting (Figure 1a). A representative single-molecule fluorescence intensity time trace (green: donor channel; red: acceptor channel) is shown in Figure 1b. Along with intensity information, the detection system also recorded photon arrival times, *i*.*e*., the time between photoexcitation and photon emission. Fluorescence lifetime decay curves were constructed for each channel, where Figure 1c shows representative curves for the FRET region of Figure 1b. Both donor and acceptor decay curves were fit using an exponential function to extract the FRET timescale. For the acceptor curve, the two components correspond to the FRET-sensitized acceptor rise time (*τ*_*rise*_) and the lifetime of the acceptor only (*τ*_*A*_). For the donor curve, the components correspond to the lifetime of the donor in the presence of the acceptor (*τ*_*DA*_). In an ideal system, *τ*_*rise*_ and *τ*_*DA*_ are equivalent.

**Figure 1:**
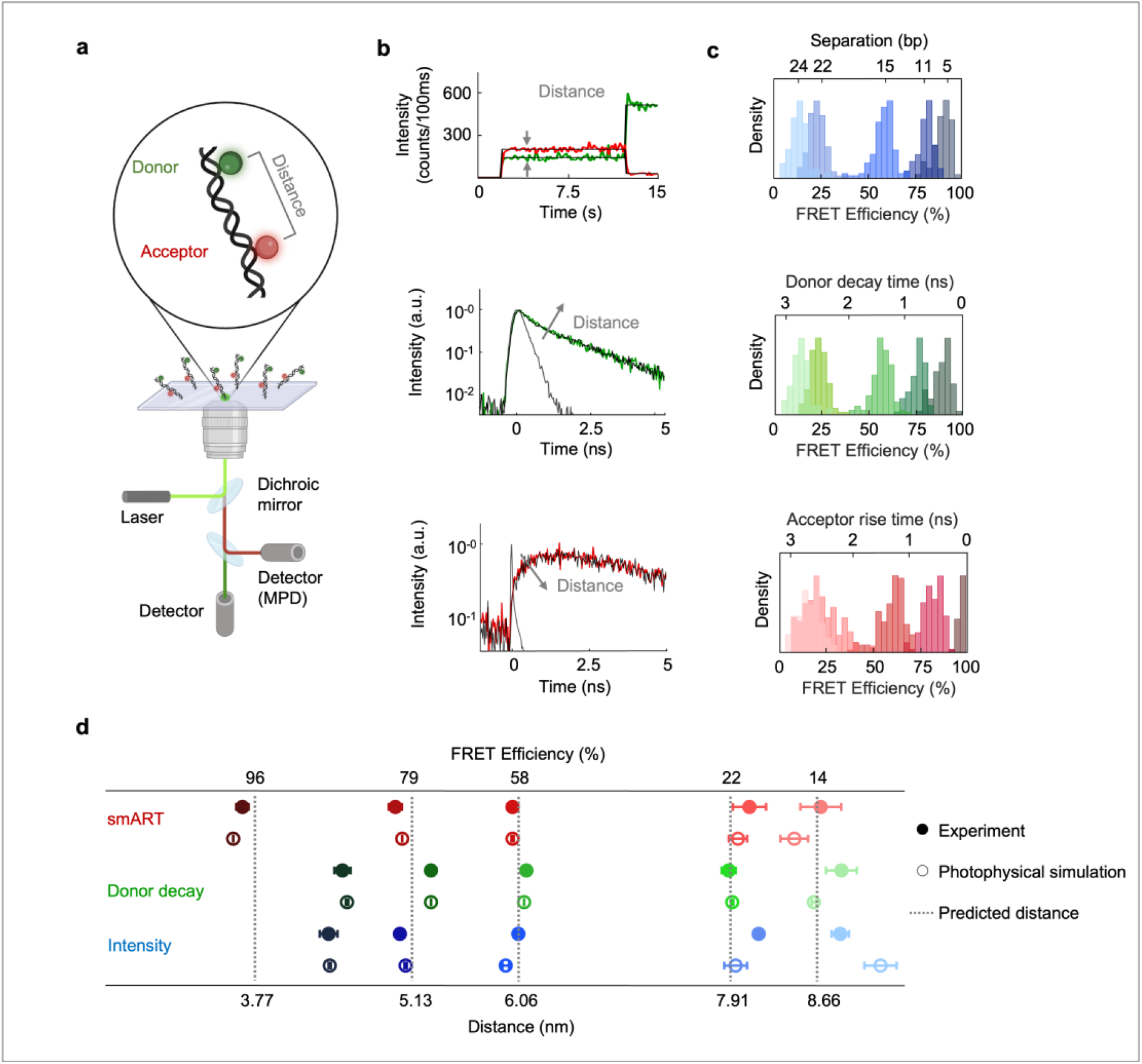
smFRET measurements on DNA rulers. **a**, Schematic of single-molecule confocal microscope. **b**, Representative single-molecule trace of fluorescence intensity from donor (green) and acceptor (red) conjugated to a DNA ruler (a, inset). Anticorrelated intensity indicates the presence of FRET. The corresponding fluorescence decay curves are donor, green; acceptor, red; fit, black. Arrows indicate the changes in the decay profile upon increasing donor-acceptor separation. The instrument response function (IRF) of the detector is shown in gray. **c**, The histogram distribution of intensity-based, donor decay, and acceptor rise time of high-high FRET, high FRET, mid FRET, low FRET, and low-low FRET. **d**, Mean FRET efficiencies and their corresponding distances for the five base pair distances reported in c. The close circles represent the experimental result. The open circles represent the simulated result. The grey dotted lines represent theoretical distances and FRET efficiencies.

To evaluate the performance of smART FRET across a range of efficiencies, DNA constructs were designed with donor-acceptor separations of 5 bps, 11 bps, 15 bps, 22 bps and 24 bps with predicted efficiencies of 96%, 79%, 58%, 22% and 14%, respectively (Supplementary Figure 1)^22^. The fluorescence emission was measured for approximately 100 individual constructs for each of the five samples. FRET efficiency was calculated from the measured data in three different ways: the relative intensities of the two channels, the donor decay time, and the acceptor rise time. For intensity-based calculations, the efficiency was determined using the following equation, 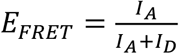 where *I*_*D*_ and *D*_*A*_ are the corrected donor and the acceptor fluorescence intensities, respectively. The efficiency was also determined using 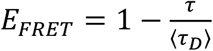, where ⟨*τ*_*D*_⟩ is the average donor lifetime and *τ* is *τ*_*DA*_ or *τ*_*rise*_ for the donor-decay and smART methods, respectively. The efficiency was converted to distance using the donor and acceptor steady-state spectra (Supplementary Figure 2).

The fluorescence emission was analyzed using all three methods to calculate the FRET values for all measured constructs. Distributions of the efficiency and distance were generated for each method (Figure 1c, Supplementary Figure 4). Figure 1d shows the mean and error of each distribution along with the expected distances for DNA duplexes (Supplementary Table 4). Using these predictions as a benchmark, the values determined using smART FRET were accurate for efficiencies of 15-95%. In contrast, the FRET efficiencies and distances calculated by donor decay and intensity were accurate around efficiencies of 50%, but showed deviations away from this region.

### Photophysical origin of improved performance of smART FRET

To investigate the photokinetic origin of the better performance of smART-based technique, we used the empirical photophysical heterogeneity from donor- and acceptor-only measurements, calculated FRET rates^23,24^, experimental signal and background levels, and Poissonian noise to numerically simulate the FRET efficiency distributions for each analysis method (Supplementary Figure 12-17). Figure 1d displays the simulated FRET efficiencies (open circles) along with the experimental (closed circles) and theoretical efficiencies (dotted vertical lines). Strong agreement between experiment and simulation was observed across all distances and for all methods.

The smART method recovers accurate values across the 15-95% range. In contrast, intensity and donor decay methods deviate from the true values, particularly in certain regimes where signal levels are low. Examination of the simulations allows us to identify the experimental parameters responsible for the deviation. In the high FRET regime (>75%), the low signal level and corresponding high signal-to-background ratio in the donor channel artifactually lower the measured efficiencies for methods that use donor emission, *i*.*e*., intensity and donor decay. The background emission appears in the spectral region surrounding the laser excitation (donor channel), yet does not have a significant contribution elsewhere (acceptor channel; Supplementary Figure 5).^22^ As a result, smART FRET is robust to these effects, and thus is designed as a nearly background-free measurement.

In the low FRET regime (<20%), the intensity method deviates from the true values due to low photon counts in the acceptor channel. That is, the donor intensity is near zero for >75% efficiencies, whereas the acceptor intensity is near zero for <20%. At these near-zero intensities, the fluorescence emission, including any changes in this emission, fall below the noise floor. These constraints lower the accuracy and limit the applicability to dynamic systems that span different FRET values.

Finally, intensity and donor decay methods calculations are subject to the inherent, underlying heterogeneity of these photophysical parameters, as observed in the donor only measurements (Supplementary Figures 3, 12). The heterogeneity is then convolved with FRET, adding noise to the recovered efficiency. In contrast, the acceptor rise time is spectroscopically separated from photophysical heterogeneity, providing a higher signal-to-noise readout for FRET determination. Essentially, the rise time directly isolates the FRET process, which leads to a higher signal-to-noise ratio than indirect methods^25^. Collectively, these advantages are reflected in the improved performance of smART FRET. The direct spectroscopic readout with a steep distance dependence over the entire nanometer-scale range allows high sensitivity, even in regimes with low signal levels.

### Single-stranded DNA-binding protein from *E. coli*

The single-strand DNA binding protein (SSB) is involved in all aspects of DNA metabolism, including replication, recombination, and repair^26,27^. SSB is a tetrameric protein that binds ssDNA^28^. Depending on the salt and free SSB concentrations, the SSB-ssDNA complex adopts different conformations, which are thought to serve different roles in genome regulation^29,30^. In high salt conditions (>200mM NaCl), approximately 65 nucleotides of ssDNA fully wrap around the tetrameric SSB^31,32^. Decreasing the salt concentration is known to cause a second SSB bind to the SSB-ssDNA complex, where each tetramer SSB occupies 35 nucleotides of the ssDNA^30,32,33^. To establish the utility of smART FRET as a versatile tool for studies of biomolecules, we investigated the architecture of single-stranded DNA (ssDNA) bound to SSB under different conditions (Figure 2).

**Figure 2:**
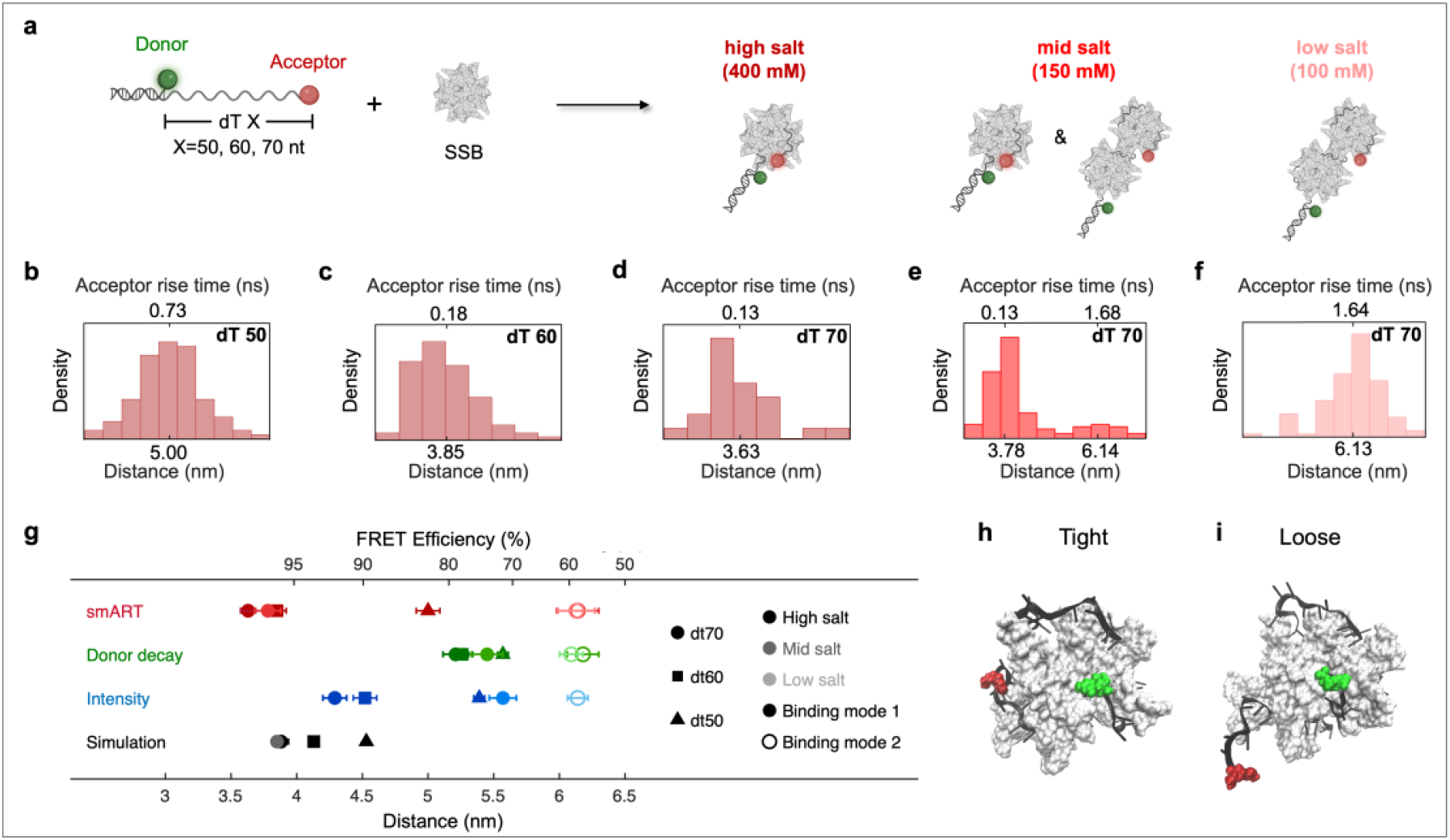
smART FRET of the single-stranded binding protein (SSB) from *E. coli*. **a**, Schematic of DNA duplex with an ssDNA overhang composed of X-nucleotide (nt) poly-dT (dT50/dT60/dT70). The inter-dye distance for the X-nt ssDNA constructs upon binding with SSB was measured in different salt conditions (400mM, 150mM, 100mM NaCl) represented by different shades of red. (**b-f)** smART FRET efficiency histograms for: **b**, dT50 binding to SSB in high salt (400mM). **c**, dT60 binding to SSB in high salt (400mM). **d**, dT70 binding to SSB in high salt (400mM). **e**, dT70 binding to SSB in mid salt (150mM). **f**, dT70 binding to SSB in low salt (100mM). **g**, Summary of the FRET results based on smART (red), donor decay (green), intensity (blue), and simulation (black). The circle represents dT70; square represents dT60; triangle represents dT50. Different salt concentrations are represented by different color shades. The close circle represents binding mode 1, and the open circle represents binding mode 2. **h**,**i;** Molecular dynamics snapshots of the ‘tight’ and ‘loose’ conformations, corresponding to distances of 3.75 nm and 4.5 nm, respectively.

We designed a partial DNA duplex with an ssDNA overhang composed of poly-dT in three different lengths (50-nucleotide poly-dT (dT50), 60-nucleotide poly-dT (dT60), 70-nucleotide poly-dT (dT70)). The donor dye (Atto 550) was conjugated to the short strand and the acceptor dye (Atto 647N) was conjugated to the long strand, generating a construct with the donor at the start of the duplex segment and the acceptor at the far end of the overhang (Figure 2a). A biotin affinity tag was introduced on the donor strand to immobilize the sample on a coverslip in a solution of 1.32 nM of SSB at high salt (400 mM NaCl).

FRET distributions were constructed for each sample using smART (Figure 2b-d), donor decay, and intensity (Supplementary Figure 8). The mean FRET efficiencies and distances for all distributions are given in Figure 2g and in Supplementary Table 6. The experimental FRET distributions showed efficiency values ranging from 73% - 81% for the dT50, 76% - 95% for the dT60, and 77% - 97% for the dT70 across the three methods of analysis. The lower FRET efficiency observed for the dT50 construct is likely because the shorter strand cannot fully encircle the SSB, whereas the high efficiencies for dT60 and dT70 suggest near-total or total encircling of the SSB. These results are consistent with previous results that SSB is fully encircled by 65 nucleotides in high salt^31,32,34^. The smART method measured distance of ∼3.6 nm for the dT70, whereas the intensity and donor decay methods measured distances of 4.3 and 5.2 nm, respectively. The difference between methods is consistent with the discrepancy observed for the DNA ruler in the high FRET regime (5 bps; Figure 1d), highlighting that the improved performance of smART FRET established in the proof-of-principle experiments is also likely present in the more complex, bimolecular SSB-ssDNA system.

### Salt dependence of the SSB-ssDNA complex

To examine how the SSB-ssDNA architecture is regulated by salt, we compared the SSB binding of the dT70 nucleotide under high salt (400 mM NaCl) to that under mid salt (150 mM NaCl) and low salt (100 mM NaCl) conditions (Figure 2a, d-f, h). In high salt, the smART FRET distribution showed a single peak with a mean at 3.63 nm distance separation between the ends of the ssDNA, corresponding to the fully encircled 65 nucleotide binding mode. In mid salt (150mM), the smART FRET efficiency distribution was bi-modal with 3.78 nm and 6.14 nm populations, corresponding to the 65 nucleotide and 35 nucleotide binding modes, respectively, similar to previous observations^32,34^. Notably, the 65 nucleotide binding mode maintained the tight binding of the ssDNA observed under high salt conditions, suggesting the ssDNA organization is robust to the local environment and the salt modulates the binding mode exclusively through the interaction between the SSB tetramers^34^. In low salt (100 mM), the smART FRET distribution showed a single peak at 6.13 nm, corresponding to the 35 nucleotide binding mode only.

The mid salt data shows that smART FRET has an improved ability to resolve bimodal distributions as compared to the other methods. The smART analysis resolved the two binding modes with a ∼2.5 nm distance separation, whereas the donor decay analysis resolved a ∼0.7 nm separation. The close spacing is likely due to the deviation of high FRET values to lower efficiencies as a result of the low signal and high background in the donor channel^35^. The intensity-based analysis failed to even resolve the bi-modal distribution at mid salt. The higher accuracy and precision from smART FRET thus enhance the visualization of these binding modes relative to traditional methods, a feature that could be invaluable in complex systems with congested conformational landscapes.

### MD simulations of SSB-ssDNA

To further understand the structural details, we performed molecular dynamics (MD) simulations on an SSB tetramer with a 70-nucleotide ssDNA bound in the 65-nucleotide binding mode (Supplementary Figure 18c-e and Table 9) under 100 mM, 150 mM, and 400mM NaCl conditions^36,37,38^. The distance between the first and last nucleotides of the ssDNA strand are plotted in Figure 2g. The distances between pairs separated by 50 or 60 nucleotides within the full 70-nucleotide ssDNA were also extracted. This resulted in a set of distances for each nucleotide separation, with each distance corresponding to a unique contact mode between the ssDNA strand and SSB. The average was calculated for the set of all extracted distances for both the dT50 and dT60 nucleotides. These values for all three residue separations are shown in Figure 2g, Supplementary Figure 18a-c and Table 9.

Overall, the averaged simulated distances recapitulated our experimental trend. The simulated distances for the dT60 and dT70 were similar to our smART FRET results (0.28 nm and 0.25 nm difference, respectively), whereas larger deviations were present for donor decay (1.13 and 1.33 nm differences, respectively) and intensity-based methods (0.39 and 0.41 nm differences, respectively). The agreement between the smART FRET and computational results supports the accuracy of the new method as well as the presence of ‘tight’ binding within the SSB-ssDNA complex. Slightly larger differences between experimental and simulated values were observed for the dT50 construct (0.47 nm) from smART FRET. The DNA duplex is present in the experimental constructs but absent in the simulations, and could block certain configurations observed in silico, leading to a mismatch between the values. Additionally, some binding positions could be more thermodynamically favorable, and so there could be uneven weighting of contact modes in the distance averages. The simulations for the dT70 under 150 mM NaCl also reported very similar distance to our experimental result with 0.07 nm difference (Figure 2h), supporting that SSB-ssDNA binding has a tight conformation under a range of salt conditions.

In both high and medium salt conditions, close proximity was observed between the termini of the bound ssDNA in both the smART FRET and simulated data, indicating that the ssDNA is held more tightly by the SSB along the full 65 nucleotide length than the other FRET methods suggest^39^. These short end-to-end distances show that SSB holds the ssDNA in a ‘tight’ conformation, as opposed to allowing the ends to detach and fluctuate in solution in a more ‘loose’ conformation. Snapshots of the simulations were extracted at distances corresponding to the predominant tight conformation (82%) and rare loose conformation (18%) (Figure 2 h-i, Supplementary Figure 19, Table 10). The tight binding observed with smART FRET may serve to protect the ssDNA during metabolism by limiting access of other reactive species in the cell^39,40,41^. Further, the persistence of this tightly bound conformation across salt concentrations suggests this strong SSB-ssDNA interaction is not directly modulated by salt, but rather the salt-dependence of SSB conformations is regulated through the interactions between SSB tetramers. Consistently, dedicated protein machinery is required to unwrap ssDNA from SSB after formation of the SSB-ssDNA complex as a part of the regulation of DNA metabolism ^42,43^. If the SSB-ssDNA complex were typically in the open conformation with the ssDNA easily disassociating, unwrapping may occur spontaneously, interfering with the tight regulation of this process^44,45^.

## Discussion

smART FRET introduces a method that leverages a spectroscopic readout separate from the photophysical noise and background emission that limits other readouts. The improved precision, accuracy and range of smART FRET establishes that this modality can be broadly used for structural and dynamical studies. The lifetime-based nature of the measurement does not currently allow widefield implementations, preventing parallelized acquisition of single-molecule measurements. Nevertheless, with recent advances in time-resolved cameras^46^, we envision that such apparatuses will be possible in the near future. Inter- and intramolecular distances are crucial experimental observables, and the high resolution newly afforded by smART FRET has the potential to reveal the interactions and reactions that underpin function in a wide range of systems.

## Methods

### Sample preparation

All DNA oligonucleotides were ordered from Integrated DNA Technologies, which synthesized and labeled the single DNA strands with dyes and carried out HPLC purification. All sequences and modifications of nucleic acids used in this study are summarized in Supplementary Table 1. Single-stranded DNA binding protein from *E*. coli was purchased from ThermoFisher Scientific. All other chemicals for buffer preparation were ordered from Sigma-Aldrich.

### DNA ruler sample

200 nM FRET samples were annealed in hybridization buffer 1 consisting of 20 mM MgCl_2_, 5 mM NaCl, 5 mM Tris, pH 7.5 for 1 hour in room temperature. Then 200 nM DNA samples were diluted to 200 pM before preparing the single-molecule sample. Immobilized single-molecule samples were prepared on a low-density biotin coverslip (MicroSurfaces, Bio 01). Samples were measured in a chamber comprised of 1X TAE buffer and oxygen scavengers (2 mM Trolox, 2.5 mM protocatechuic acid and 25 nM protocatechuic-3,4-dioxygenase).

### SSB–ssDNA complex sample

100 nM DNA samples were annealed in hybridization buffer 2 which consists of 20 mM Tris, 400 mM NaCl, 10 mM MgCl_2_, pH 7.4 for 1 hour at room temperature. Then 0.66 uM of SSB was added to the 100 nM DNA samples, incubated for 30 minutes at room temperature. Immobilized single-molecule samples, with final concentrations of 200 pM DNA and 1.32 nM SSB, were prepared on a low-density biotin coverslip (MicroSurfaces, Bio 01). Samples were measured in a chamber comprised of hybridization buffer 2 and oxygen scavengers (2 mM Trolox, 2.5 mM protocatechuic acid and 25 nM protocatechuic-3,4-dioxygenase).

## Experimental setup

**FRET experiments** were carried out on a home-built confocal microscope with two separate spectral detection channels for donor and acceptor emission (Figure 1A) as described previously ^47^. Excitation light was generated using a Ti-sapphire laser (Coherent, Vitara-S; λc = 800 nm, 70 nm bandwidth, 20 fs pulse duration, 80 MHz repetition rate) focused into a nonlinear photonic crystal fibre (NKT Photonics, FemtoWhite 800) and then filtered to 550 nm (Omega, RPB 540/560). An oil-immersion objective (Olympus; UPLSAPO100XO, NA = 1.4) was used to focus the excitation to the sample and collect the resultant fluorescence. The fluorescence was passed through a dichroic filter (Laser 2000, SP01-561RU) with donor bandpass filter (Thorlabs, FB600-40) and acceptor bandpass filter (Chroma, ET700/75m). Fluorescence emission was detected by two kinds of photon-counting avalanche photodiodes, Excelitas, SPCM-AQRH-15 for the donor channel and Micro Photon Devices, PDM for the acceptor channel. The instrument response function (IRF) for the donor detector was measured to be ∼380 ps (full width at half-maximum) and for acceptor detector was measured to be ∼80 ps using scatter off a glass coverslip. Photon arrival times were recorded by a time-correlated single-photon counting module (Picoquant, PicoHarp 300). A 5 μm × 5 μm area of a coverslip with immobilized samples was initially scanned. Diffraction limited and spatially separated single molecule spots were then probed individually by unblocking the laser beam to record fluorescence until photo-bleaching. The laser power for the experiments was ∼3 μW. Florescence emission was binned at 100-ms resolution to generate fluorescence intensity traces for both the donor and acceptor channels. Traces with a single photobleaching step for the donor and acceptor were considered for further analysis.

**Acceptor-only experiments** were performed on another home built confocal microscope as described in ref^48^. In brief, a tunable fiber laser (FemtoFiber pro, Toptica; ∼100 fs pulse duration, 610 nm, ∼4 nm bandwidth, 80 MHz repetition rate) was used as the excitation source. The excitation was focused by an oil-immersion objective (UPLSAPO100XO, Olympus, NA 1.4) onto the samples immobilized on a coverslip. The coverslip was mounted on a piezostage controlled by a home-written Labview-based software. The emission of the sample was collected through the same objective and separated from the excitation by using band pass filter (Chroma, ET700/75m). The average power of excitation was ∼1 μW. Emission was detected by a single photon counting avalanche photodiode (SPCM-AQRH, Excelitas Technologies). A time-correlated single-photon counting (TCSPC) module (Time Tagger 20, Swabian Instruments) was used to record the arrival times of the photons.

## Data analysis

smFRET investigation performed on confocal microscopy benefits from sub-millisecond time resolution and can achieve near-shot noise limited precision. Florescence emission was binned at 100-ms resolution to generate fluorescence intensity traces for both the donor and acceptor channels. Traces with a single photobleaching step for the donor and acceptor were considered for further analysis. Regions of constant intensity in the traces were identified by a change-point algorithm^49^. All traces were background corrected with the intensity detected after the fluorophores photobleached. The kinetic traces were fit using maximum likelihood estimation (MLE). Fluorescence intensity and lifetime were analyzed as described previously^50^. Both donor and acceptor decay curves were fit by a two-exponential function convolved with the instrument response function (IRF). The fit was performed using maximum likelihood estimation (MLE). The donor and acceptor intensity decay are defined by the following equations:

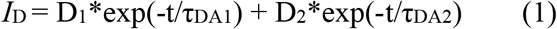

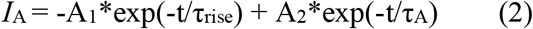

Where *I*_D_ and *I*_A_ are the experimentally observed intensity of donor and acceptor; D_1_, D_2_, A_1_, A_2_ are the amplitudes of the lifetime components; *τ*_DA1_ and *τ*_DA2_ are two components of donor decay times, the average data of *τ*_1_ and *τ*_2_ were used as FRET-quenched donor decay time *τ*_DA_ which is used for the efficiency and distance calculation; *τ*_rise_ is FRET-sensitized acceptor rise time; *τ*_A_ is the acceptor-only lifetime with unpresented donor.

The acceptor rise time, donor decay time and intensity ratio can be used to quantify energy transfer efficiency and then converted to center-to-center distance between donor and acceptor dyes. The FRET efficiency is defined by:

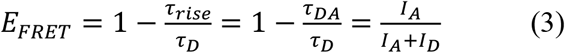

Intensity-based determination of FRET efficiencies requires consideration of the following correction factors: α, a factor for spectral cross-talk arising from donor fluorescence leakage in the acceptor channel; δ, a factor for direct excitation of the acceptor with the donor laser; and a detection correction factor (γ). The spectral cross-talk from donor and direct excitation of the acceptor were calculated from the filter spectra, and the detection correction factor was calculated from the detector efficiency and fluorophore quantum yield. The FRET efficiency defined by intensity was corrected in this work using the calculated factors and the following equation:

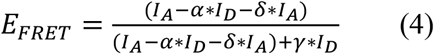

The distance was calculated from the FRET efficiency using:

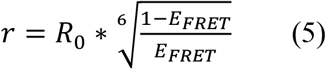

Where *E*_FRET_ is FRET efficiency; *R*_0_ is the Förster distance (the donor-acceptor distance for which FRET is 50 % efficient); *r* is the donor-acceptor distance.

A Förster distance of *R*_0_ = 6.4 nm was calculated for the dyes used in this study using the following equation:

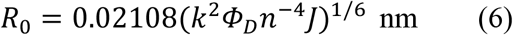

The orientation factor κ^2^ was taken as 2/3 because of random orientation of donor and acceptor during the FRET time. The refractive index was *n* = 1.35 (aqueous solution). The Atto550 quantum yield was Φ_D_ = 0.80. The overlap integral *J* was calculated by the following equation:

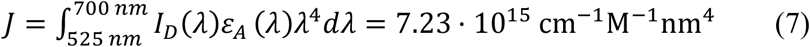

*I*_D_(λ) is the emission intensity from the area-normalized (to unity) emission spectrum of donor and ε_A_(λ) is the molar absorptivity of the acceptor.

