## Supplementary for "Single-molecule acceptor rise time (smART) FRET for nanoscale distance sensitivity"

#### Contents

#### Part 1: Experiment data

1. Supplementary Figure 1: Theoretical FRET Efficiency and distance prediction of DNA ruler

2. Supplementary Figure 2: Photophysical properties of Atto550 and Atto647N, and $R_0$ calculation

3. Supplementary Figure 3: Distribution of donor-only and acceptor-only lifetime of DNA ruler

4. Supplementary Figure 4: Histograms of lifetimes, FRET efficiencies, and distances for the DNA rulers

5. Supplementary Figure 5: Representative single molecule fluorescence traces of DNA ruler

6. Supplementary Figure 6: Representative single molecule fluorescence lifetime decays of DNA ruler

7. Supplementary Figure 7: Distribution of donor-only and acceptor-only lifetime of SSB-DNA system

8. Supplementary Figure 8: Distribution of SSB-DNA binding system with an ssDNA overhang composed of dT50, dT60 and dT70 under high salt condition

9. Supplementary Figure 9: Representative single molecule fluorescence traces of SSB-DNA system

10. Supplementary Figure 10: Representative single molecule fluorescence lifetime decays of SSB-DNA system

11. Supplementary Figure 11: Distribution of SSB-DNA binding system of dT70 ssDNA overhang under mid and low salt conditions

12. Supplementary Table 1: DNA oligonucleotides used in the study

13. Supplementary Table 2: Dye parameters for the AV simulations with AV3-model.

14. Supplementary Table 3: Lifetime of donor-only and acceptor-only of DNA ruler

15. Supplementary Table 4: FRET results of DNA ruler

16. Supplementary Table 5: Lifetime of donor-only and acceptor-only of SSB-DNA system

17. Supplementary Table 6: FRET results of SSB-DNA system with an ssDNA overhang composed of dT50, dT60 and dT70 under high salt condition

18. Supplementary Table 7: FRET results of SSB-DNA binding system of dT70 ssDNA overhang under mid and low salt conditions

#### Part 2: Numerical simulation of smFRET system of DNA ruler

1. Supplementary Figure 12: Distribution of donor-only intensity and γ factor

2. Supplementary Figure 13: Algorithm to simulate intensity-based smFRET efficiency

3. Supplementary Figure 14: Algorithm to simulate donor lifetime decay-based smFRET efficiency

4. Supplementary Figure 15: Distribution of the donor decay lifetimes of the experiments and the simulations

5. Supplementary Figure 16: Algorithm to simulate acceptor rise time-based smFRET efficiency

6. Supplementary Figure 17: Distribution of the acceptor rise time of the experiments and the simulations

7. Supplementary Table 8: The FRET efficiency results of experiments and the simulations.

#### Part 3: Molecular dynamics simulation of SSB-DNA

1. Supplementary Figure 18: Traces of SSB-DNA molecular dynamic.

2. Supplementary Figure 19: Bi-modal fitting of SSB-DNA (dT70) molecular dynamic simulation in 400mM NaCl.

3. Supplementary Table 9: Summary of SSB-DNA molecular dynamic simulation results.

4. Supplementary Table 10: Summary of bi-modal fitting of SSB-DNA (dT70) molecular dynamic simulation in 400mM NaCl.

### Part 1: Experiment data

#### Supplementary Figure 1: Theoretical FRET efficiency and distance prediction of DNA ruler

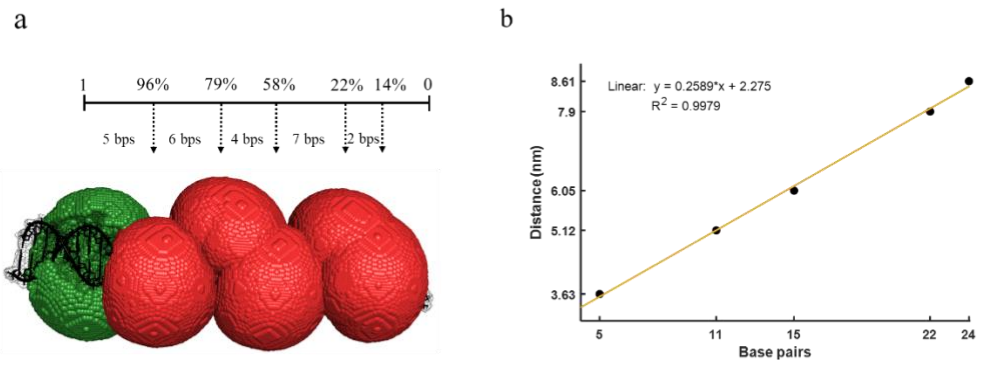


**a,** The theoretical FRET efficiency of 5 bps, 11 bps, 15 bps, 22 bps and 24bps separation. **b,** The linear calibration curve of the theoretical distance of 5 bps, 11 bps, 15 bps, 22 bps and 24bps separation.

Double-stranded DNA was labeled with Atto550 and Atto647N at 5 bps, 11 bps, 15 bps, 22 bps and 24bps separation referred to as “high-high FRET”, “high FRET”, “mid FRET”, “low FRET” and “low-low FRET” samples, respectively. The model for the double-stranded DNA was generated using the Avogadro software[1]. The dye molecules were modeled as ellipsoids (approximated by three radii; AV3-Model) and AVs were generated using the FPS software[2]. For the efficiency calculation at a given distance, the Förster radius specific to each dye pair was used. The labelling position was the C7 of the thymine (the C-atom of the thymine’s methyl group). All mean geometric dyes parameters are shown in Supplementary Table 2. Further used parameters are: Boundary tolerance 0.5, accessible volume grid (rel.) 0.2; Min. grid [Å] 0.4, Search nodes: 3 and E samples: 200.

**Supplementary Figure 2: Photophysical properties of Atto550 and Atto647N, and $R_0$ calculation**

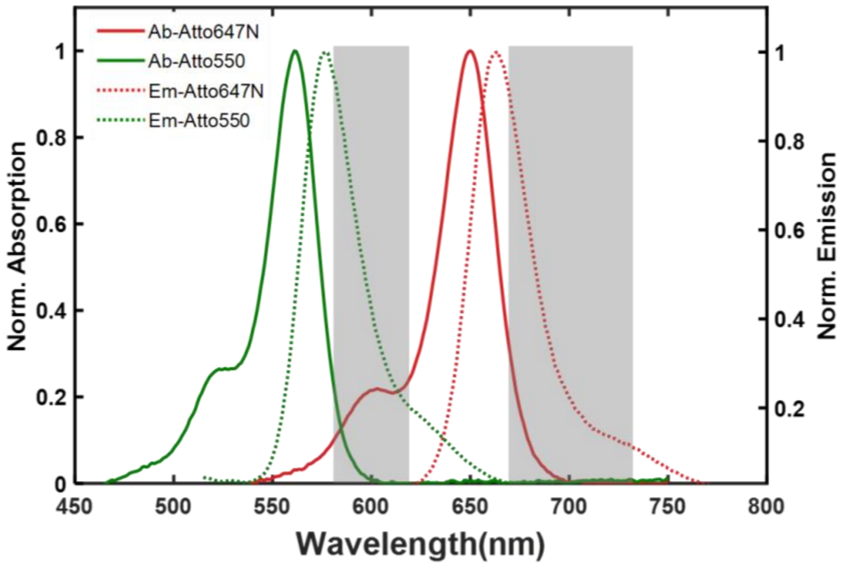


Absorbance spectra Atto550-DNA (green) and Atto647N-DNA (red). PL emission spectra of Atto550-DNA (green dash) and Atto647N-DNA (red dash). Optical bandpass filter transmission spectra are shown in gray.

**Supplementary Figure 3: Distribution of donor-only and acceptor-only lifetime of DNA ruler**

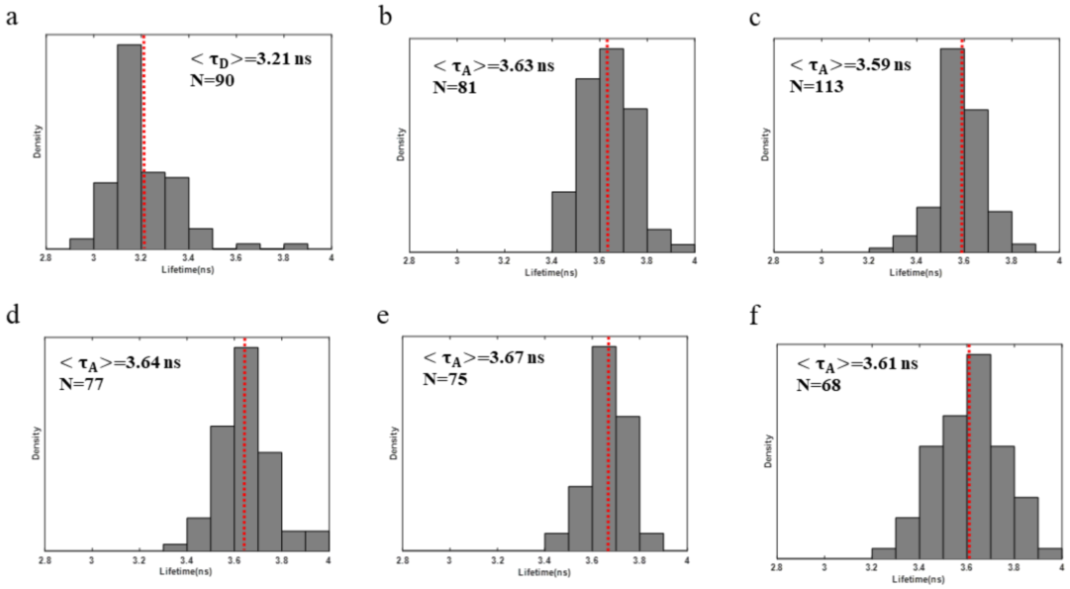


**a,** Histogram distribution of donor-only lifetime. **b,** Histogram distribution of acceptor-only (high-high) lifetime. **c,** Histogram distribution of acceptor-only (high) lifetime. **d,** Histogram distribution of acceptor-only (mid) lifetime. **e,** Histogram distribution of acceptor-only (low) lifetime. **f,** Histogram distribution of acceptor-only (low-low) lifetime. $<\tau>$ is the average of the single molecule lifetimes. N is the number of single molecules.

**Supplementary Figure 4: Histograms of lifetimes, FRET efficiencies, and distances for the DNA rulers**

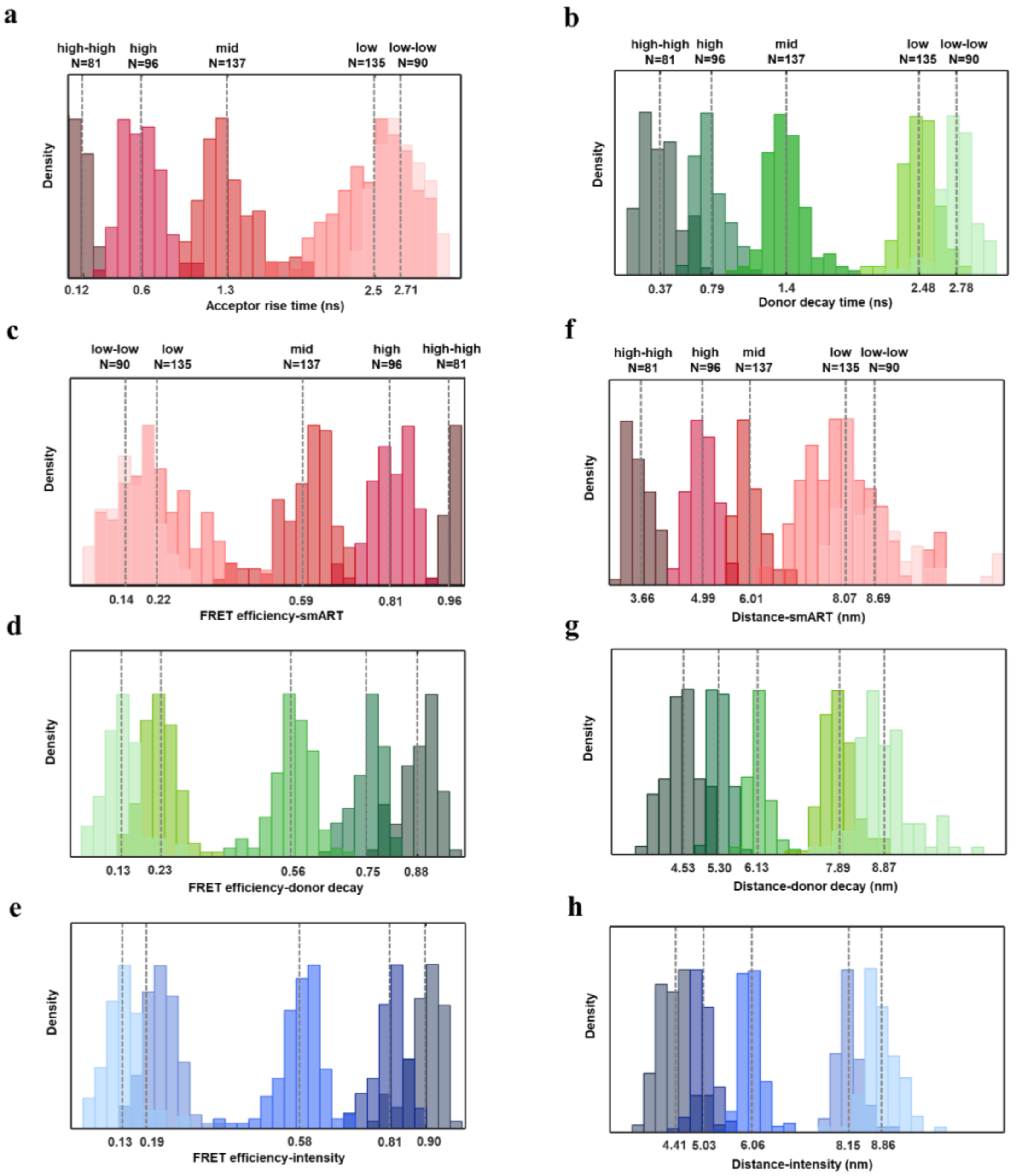


**a,** Histogram distribution of FRET acceptor rise time. **b,** Histogram distribution of FRET donor decay time. **c,** Histogram distribution of FRET efficiency based on acceptor rise time. **d,** Histogram distribution of FRET efficiency based on donor decay time. **e,** Histogram distribution of FRET efficiency based on intensity. **f,** Histogram distribution of distance based on acceptor rise time. **g,** Histogram distribution of distance based on donor decay time. **h,** Histogram distribution of distance based on intensity. high-high FRET, high FRET, mid FRET, low FRET, and low-low FRET in different shade; N is the number of the single molecules.

**Supplementary Figure 5: Representative single molecule fluorescence traces of DNA ruler**

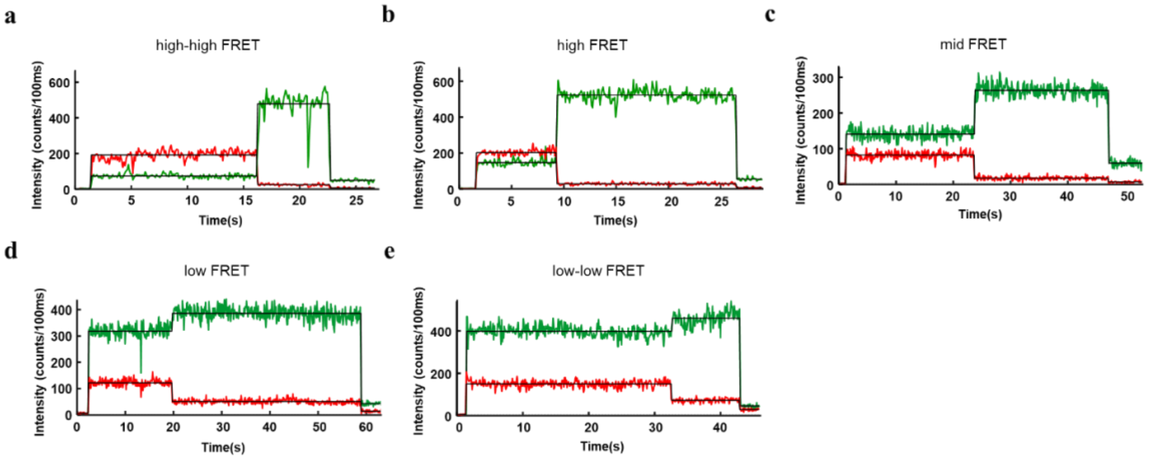


Representative single-molecule fluorescence traces of the DNA rulers with donor emission in green and acceptor emission in red. The traces shown are from DNA rulers designed for **a,** high-high FRET; **b,** high FRET; **c,** mid FRET; **d,** low FRET; and **e,** low-low FRET. The start of fluorescence emission corresponds to the laser turn on, the step-wise increase in donor emission and decrease in acceptor emission corresponds to acceptor photobleaching, and the step-wise decrease in emission in both channels corresponds to donor photobleaching.

**Supplementary Figure 6: Representative single molecule fluorescence lifetime decays of DNA ruler**

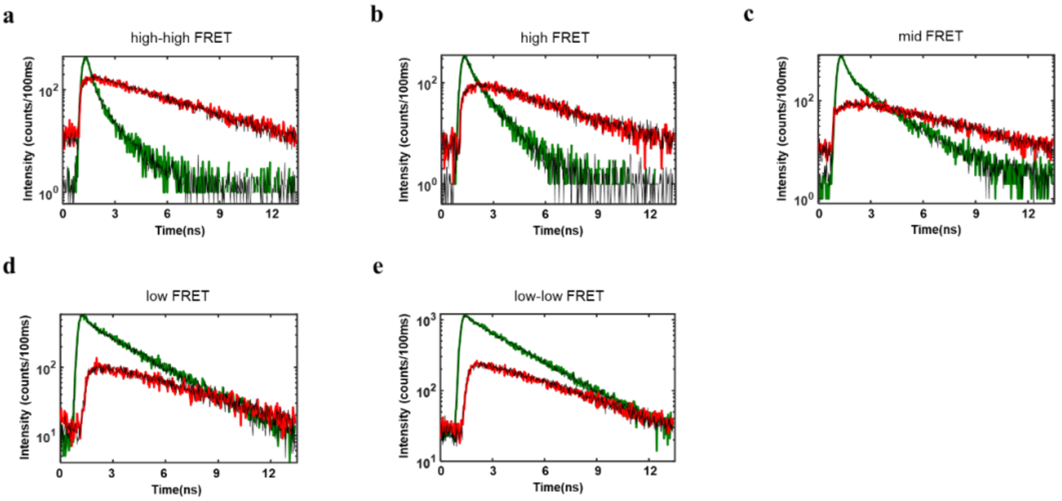


Representative single-molecule fluorescence lifetime decays of the DNA rulers with donor lifetime decay in green and acceptor lifetime decay in red. The lifetime decays shown are from DNA rulers designed for **a,** high-high FRET; **b,** high FRET; **c,** mid FRET; **d,** low FRET; and **e,** low-low FRET.

**Supplementary Figure 7: Distribution of donor-only and acceptor-only lifetime of SSB-DNA system**

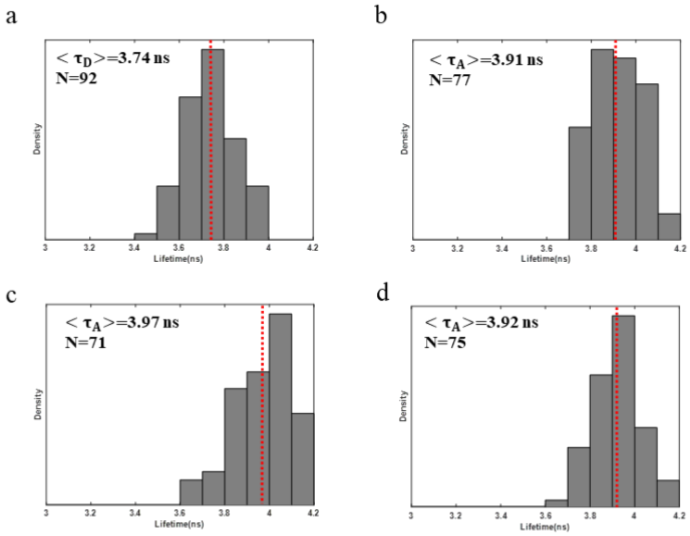


**a,** Histogram distribution of donor-only lifetime. **b,** Histogram distribution of acceptor-only (dT50) lifetime. **c,** Histogram distribution of acceptor-only (dT60) lifetime. **d,** Histogram distribution of acceptor-only (dT70) lifetime. $< \tau >$ is the average of the single molecule lifetimes. N is the number of single molecules.

**Supplementary Figure 8: Distribution of SSB-DNA binding system with an ssDNA overhang composed of dT50, dT60 and dT70 under high salt condition**

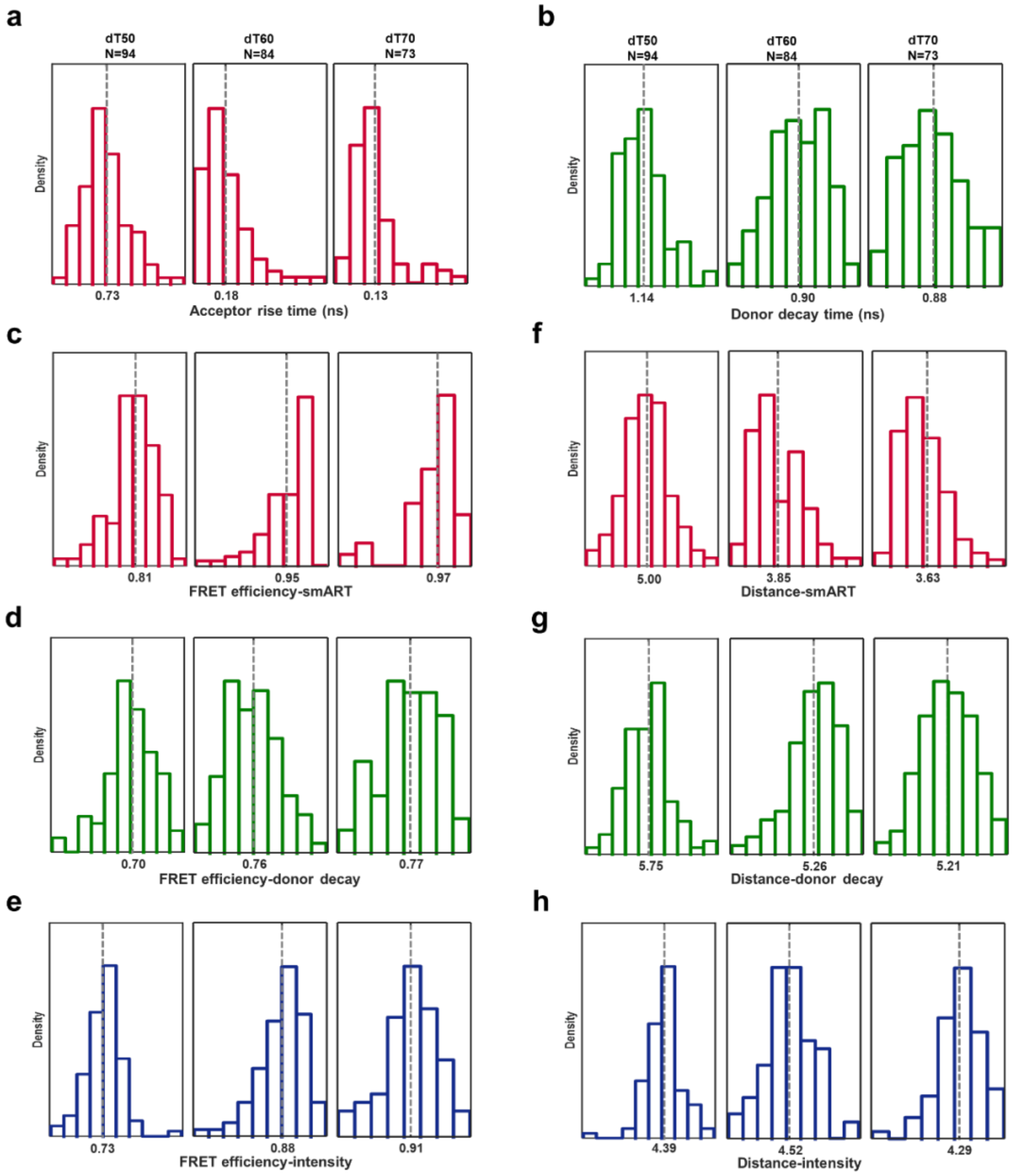


**a,** Histogram distribution of FRET acceptor rise time of dT50, dT60 and dT70. **b,** Histogram distribution of FRET donor decay time of dT50, dT60 and dT70. **c,** Histogram distribution of FRET efficiency based on acceptor rise time of dT50, dT60 and dT70. **d,** Histogram distribution of FRET efficiency based on donor decay time of dT50, dT60 and dT70. **e,** Histogram distribution of FRET efficiency based on intensity of dT50, dT60 and dT70. **f,** Histogram distribution of distance based on acceptor rise time of dT50, dT60 and dT70. **g,** Histogram distribution of distance based on donor decay time of dT50, dT60 and dT70. **h,** Histogram distribution of distance based on intensity of dT50, dT60 and dT70. N is the number of the single molecules.

**Supplementary Figure 9: Representative single molecule fluorescence traces of SSB-DNA system**

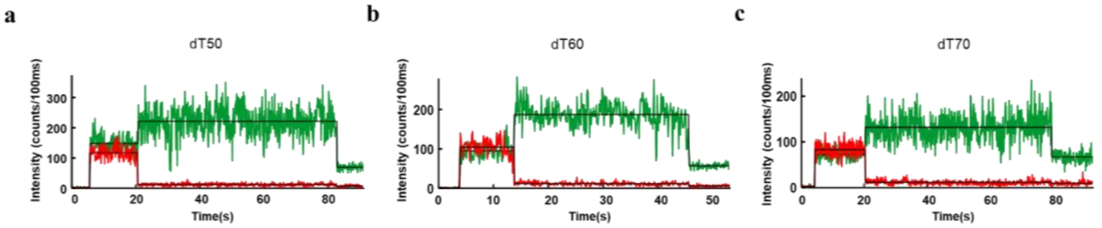


Representative single-molecule fluorescence traces of the SSB-DNA system with donor emission in green and acceptor emission in red. The traces shown are from SSB-DNA system designed for **a,** dT50; **b,** dT60; and **c,** dT70. The start of fluorescence emission corresponds to the laser turn on, the step-wise increase in donor emission and decrease in acceptor emission corresponds to acceptor photobleaching, and the step-wise decrease in emission in both channels corresponds to donor photobleaching.

**Supplementary Figure 10: Representative single molecule fluorescence lifetime decays of SSB-DNA system**

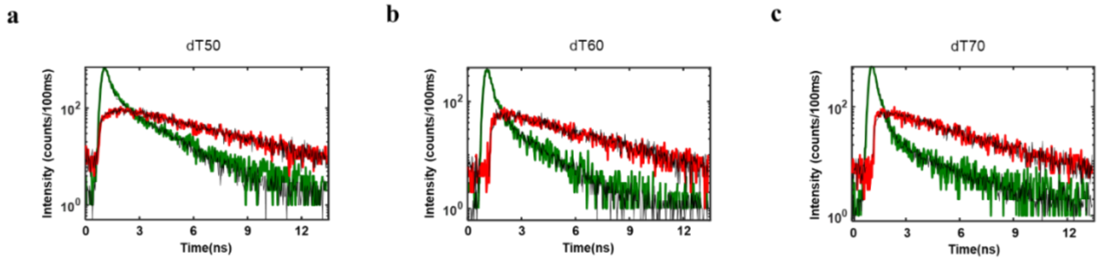


Representative single-molecule fluorescence lifetime decays of the SSB-DNA system with donor lifetime decay in green and acceptor lifetime decay in red. The lifetime decays shown are from SSB-DNA system designed for **a,** dT50; **b,** dT60; **c,** dT70.

**Supplementary Figure 11: Distribution of SSB-DNA binding system of dT70 ssDNA overhang under mid and low salt conditions**

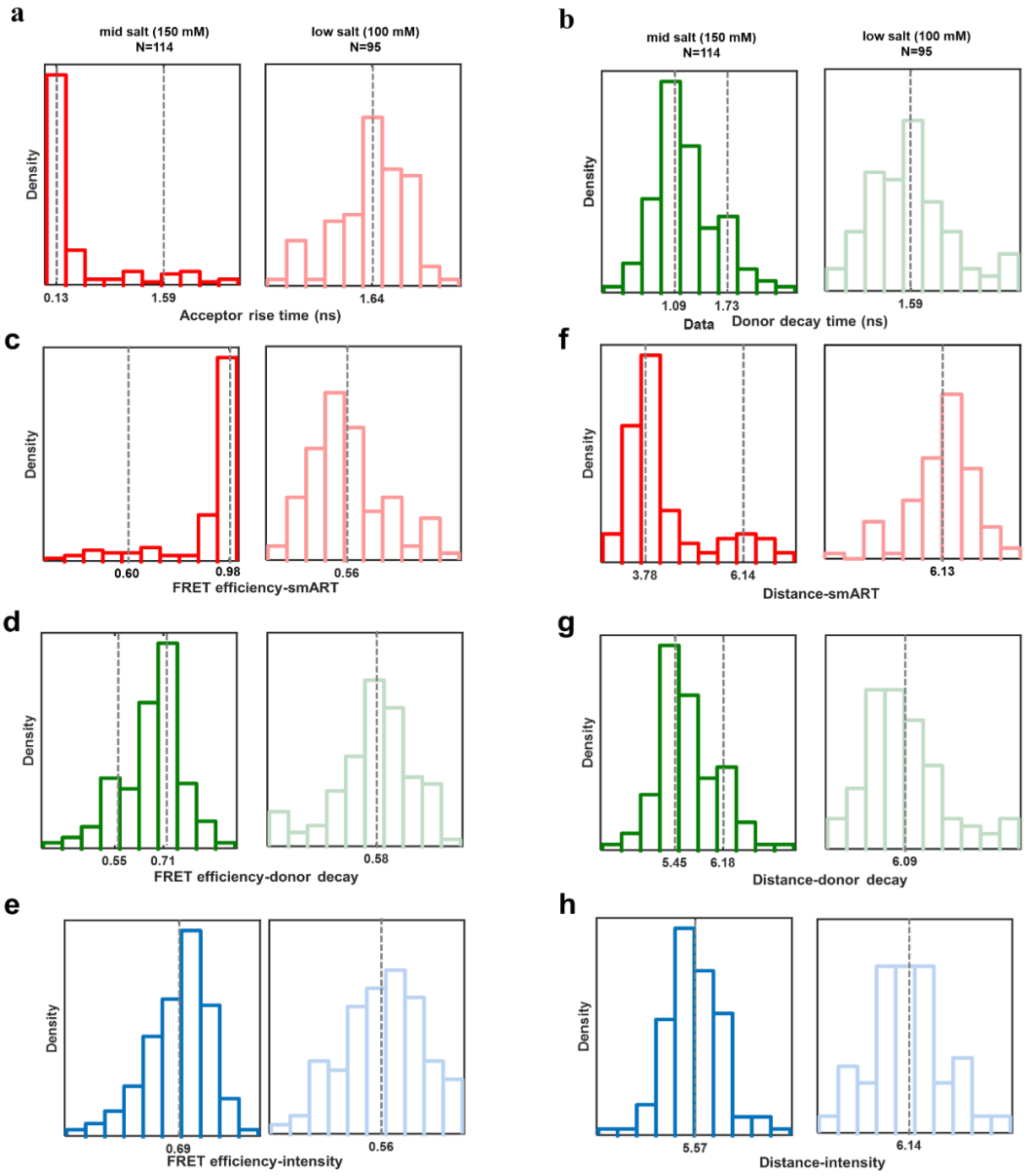


**a,** Histogram distribution of FRET acceptor rise time of dT70 under mid salt (150 mM) and low salt (100 mM) condition. **b,** Histogram distribution of FRET donor decay time of dT70 under mid salt (150 mM) and low salt (100 mM) condition. **c,** Histogram distribution of FRET efficiency based on acceptor rise time of dT70 under mid salt (150 mM) and low salt (100 mM) condition. **d,** Histogram distribution of FRET efficiency based on donor decay time of dT70 under mid salt (150 mM) and low salt (100 mM) condition. **e,** Histogram distribution of FRET efficiency based on intensity of dT70 under mid salt (150 mM) and low salt (100 mM) condition. **f,** Histogram distribution of distance based on acceptor rise time of dT70 under mid salt (150 mM) and low salt (100 mM) condition. **g,** Histogram distribution of distance based on donor decay time of dT70 under mid salt (150 mM) and low salt (100 mM) condition. **h,** Histogram distribution of distance based on intensity of dT70 under mid salt (150 mM) and low salt (100 mM) condition. N is the number of the single molecules.

#### Supplementary Table 1: DNA oligonucleotides used in the study

| Name | Sequence 5'-3' | Base position (Linker) modification |
|---|---|---|
| Donor sequence (DNA model) | CCAGACAAACACTCAAACAAACTCGACACTTTCAGCTC | dT 31(C6-Atto550), 5'-biotin |
| Acceptor sequence-high-high FRET | GAGCTGAAAGTGTCGAGTTTGTTTGAGTGTTTGTCTGG | dT 13(C6-Atto647N) |
| Acceptor sequence-high FRET | GAGCTGAAAGTGTCGAGTTTGTTTGAGTGTTTGTCTGG | dT 19(C6-Atto647N) |
| Acceptor sequence-mid FRET | GAGCTGAAAGTGTCGAGTTTGTTTGAGTGTTTGTCTGG | dT 23(C6-Atto647N) |
| Acceptor sequence-low FRET | GAGCTGAAAGTGTCGAGTTTGTTTGAGTGTTTGTCTGG | dT 30(C6-Atto647N) |
| Acceptor sequence-low-low FRET | GAGCTGAAAGTGTCGAGTTTGTTTGAGTGTTTGTCTGG | dT 32(C6-Atto647N) |
| Donor sequence (protein model) | TCGCTGCCGACTCGAGATCT | 5'-C6-Atto550, 3'-biotin |
| Acceptor sequence-dT50 | AGATCTCGAGTCGGCAGCGA(T) n, n=50 | 3'-C6-Atto647N |
| Acceptor sequence-dT60 | AGATCTCGAGTCGGCAGCGA(T) n, n=60 | 3'-C6-Atto647N |
| Acceptor sequence-dT70 | AGATCTCGAGTCGGCAGCGA(T) n, n=70 | 3'-C6-Atto647N |

#### Supplementary Table 2: Dye parameters for the AV simulations with AV3-model.

| | linker length [Å] | linker width [Å] | R1 [Å] | R2 [Å] | R3 [Å] |
|---|---|---|---|---|---|
| dT-C6-Atto550 | 20.5 | 4.5 | 7.8 | 4.5 | 1.5 |
| dT-C6-Atto647N | 20.5 | 4.5 | 7.15 | 4.5 | 1.5 |

#### Supplementary Table 3: Lifetime of donor-only and acceptor-only of DNA ruler

| Lifetime/ns Error (bootstrap) | $\tau_D$ | $\tau_A$ | | | | |
|---|---|---|---|---|---|---|
| | Donor-only | Acceptor-only | | | | |
| | | high-high | high | mid | low | low-low |
| | 3.21 | 3.63 | 3.59 | 3.64 | 3.67 | 3.67 |
| | 0.028 | 0.022 | 0.017 | 0.025 | 0.019 | 0.033 |

**Supplementary Table 4: FRET results of DNA ruler**

| Value<br>Error (bootstrap) | high-high FRET | high FRET | mid FRET | low FRET | low-low FRET |
|---|---|---|---|---|---|
| $\tau_{rise}$ (ns) | 0.11<br>0.011 | 0.60<br>0.033 | 1.30<br>0.036 | 2.50<br>0.053 | 2.71<br>0.041 |
| $E_{FRET}$ | 0.96<br>0.003 | 0.81<br>0.010 | 0.59<br>0.012 | 0.22<br>0.016 | 0.14<br>0.013 |
| Distance (nm) | 3.66<br>0.054 | 4.99<br>0.057 | 6.01<br>0.047 | 8.07<br>0.144 | 8.69<br>0.177 |
| $\tau_{DA}$ (ns) | 0.37<br>0.031 | 0.79<br>0.025 | 1.40<br>0.025 | 2.48<br>0.023 | 2.78<br>0.030 |
| $E_{FRET}$ | 0.88<br>0.011 | 0.75<br>0.008 | 0.56<br>0.009 | 0.23<br>0.008 | 0.13<br>0.009 |
| Distance (nm) | 4.53<br>0.070 | 5.30<br>0.038 | 6.13<br>0.034 | 7.89<br>0.061 | 8.87<br>0.134 |
| Intensity-$E_{FRET}$ | 0.90<br>0.010 | 0.81<br>0.008 | 0.58<br>0.009 | 0.19<br>0.005 | 0.13<br>0.006 |
| Intensity-Distance (nm) | 4.41<br>0.077 | 5.03<br>0.039 | 6.06<br>0.041 | 8.15<br>0.045 | 8.86<br>0.075 |

**Supplementary Table 5: Lifetime of donor-only and acceptor-only of SSB-DNA system**

| Lifetime(ns)<br>Error (bootstrap) | $\tau_D$ | $\tau_A$ | | |
|---|---|---|---|---|
| | Donor-only | Acceptor-only | | |
| | | dT50 | dT60 | dT70 |
| | 3.74<br>0.024 | 3.91<br>0.024 | 3.97<br>0.028 | 3.92<br>0.024 |

**Supplementary Table 6: FRET results of SSB-DNA binding system with an ssDNA overhang composed of dT50, dT60 and dT70 under high salt condition**

| Value<br>Error (bootstrap) | FRET-dT50 | FRET-dT60 | FRET-dT70 |
|---|---|---|---|
| $\tau_{rise}$ (ns) | 0.73<br>0.060 | 0.18<br>0.020 | 0.13<br>0.014 |
| $E_{FRET}$ | 0.81<br>0.016 | 0.95<br>0.006 | 0.97<br>0.004 |
| Distance (nm) | 5.00<br>0.090 | 3.85<br>0.070 | 3.63<br>0.056 |
| $\tau_{DA}$ (ns) | 1.14<br>0.033 | 0.90<br>0.056 | 0.88<br>0.074 |
| $E_{FRET}$ | 0.70<br>0.009 | 0.76<br>0.016 | 0.77<br>0.019 |
| Distance (nm) | 5.57<br>0.041 | 5.26<br>0.081 | 5.21<br>0.096 |
| Intensity-$E_{FRET}$ | 0.73<br>0.009 | 0.88<br>0.013 | 0.91<br>0.009 |
| Distance (nm) | 5.39<br>0.042 | 4.52<br>0.089 | 4.29<br>0.090 |

**Supplementary Table 7: FRET results of SSB-DNA binding system of dT70 ssDNA overhang under mid and low salt conditions**

<table>
<tr><th>Value<br>Error (bootstrap)</th><th>mid salt mode 1</th><th>mid salt mode 2</th><th>low salt</th></tr>
<tr><td>$\tau_{rise}$ (ns)</td><td>0.13<br>0.037</td><td>1.59<br>0.295</td><td>1.64<br>0.113</td></tr>
<tr><td>$E_{FRET}$</td><td>0.98<br>0.010</td><td>0.60<br>0.024</td><td>0.56<br>0.028</td></tr>
<tr><td>Distance (nm)</td><td>3.78<br>0.100</td><td>6.14<br>0.160</td><td>6.13<br>0.140</td></tr>
<tr><td>$\tau_{DA}$ (ns)</td><td>1.09<br>0.060</td><td>1.73<br>0.005</td><td>1.59<br>0.077</td></tr>
<tr><td>$E_{FRET}$</td><td>0.71<br>0.019</td><td>0.55<br>0.074</td><td>0.58<br>0.020</td></tr>
<tr><td>Distance (nm)</td><td>5.45<br>0.120</td><td>6.18<br>0.120</td><td>6.09<br>0.088</td></tr>
<tr><td>Intensity-$E_{FRET}$</td><td colspan="2">0.69<br>0.022</td><td>0.56<br>0.019</td></tr>
<tr><td>Distance (nm)</td><td colspan="2">5.57<br>0.103</td><td>6.14<br>0.077</td></tr>
</table>

#### Part 2: Numerical simulation of smFRET system of DNA ruler

##### Intensity-based FRET

To simulate the distribution of the single-molecule intensity-based FRET efficiency, we need to model donor and acceptor fluorescence intensity. This can be done by invoking the relationship of fluorescence intensity ($I$) to the radiative and non-radiative rate constants:

$$I_{D0} \propto \frac{k_r}{k_r+k_{nr}} = p * \frac{k_r}{k_r+k_{nr}} \quad \text{(Eqn. 1)}$$

where, $I_{D0}$ is the donor-only fluorescence intensity after removing the background. $k_r$ and $k_{nr}$ are the rate-constants for radiative and non-radiative decays. $p$ is a constant which depends on the donor extinction coefficients, illumination intensity and the detection efficiency of the microscope. Now, fluorescence lifetime ($\tau_D$) and quantum yield ($\phi_D$) of the donor is related by the following ways:

$$\tau_D = \frac{1}{k_r+k_{nr}} \quad \text{(Eqn. 2)}$$

$$\phi_D = \frac{k_r}{k_r+k_{nr}} \quad \text{(Eqn. 3)}$$

From the distribution of donor-only lifetime and the value quantum yield of the donor ($\phi_D = 0.8$), from the Eqn. 2 and Eqn. 3, we can obtain a distribution of $k_r$ and $k_{nr}$ for the donor. From the experimentally observed donor-only intensity distribution, the parameter $p$ can be obtained.

**Supplementary Figure 12: Distribution of donor-only intensity and γ factor**

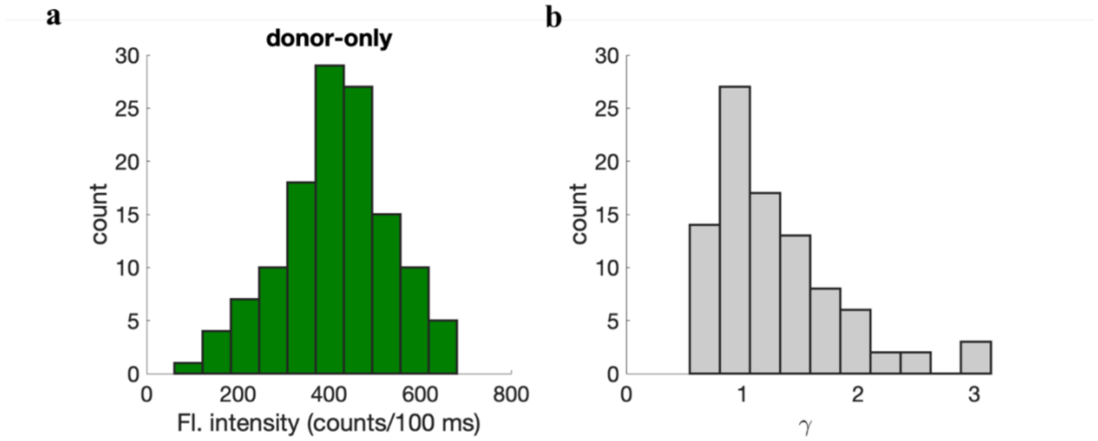


**a,** Distribution of the fluorescence intensity of the donor-only sample. **b,** factor those are used to simulate intensity-based FRET efficiencies.

In the presence of acceptor, due to the occurrence of FRET, the fluorescence intensity of the donor is reduced. This can be modeled by adding a non-radiative pathway accounting for FRET.

$$I_{DA} = p * \frac{k_r}{k_r+k_{nr}+k_t} + bg_D \quad \text{(Eqn. 4)}$$

where, $I_{DA}$ is the donor fluorescence intensity in the presence of acceptor and $k_t$ is the FRET rate. $bg_D$ is the background in the donor channel. Please note that background is added as the $p$ is calculated after the background removal of the donor-only fluorescence intensity.

**Supplementary Figure 13: Algorithm to simulate intensity-based smFRET efficiency**

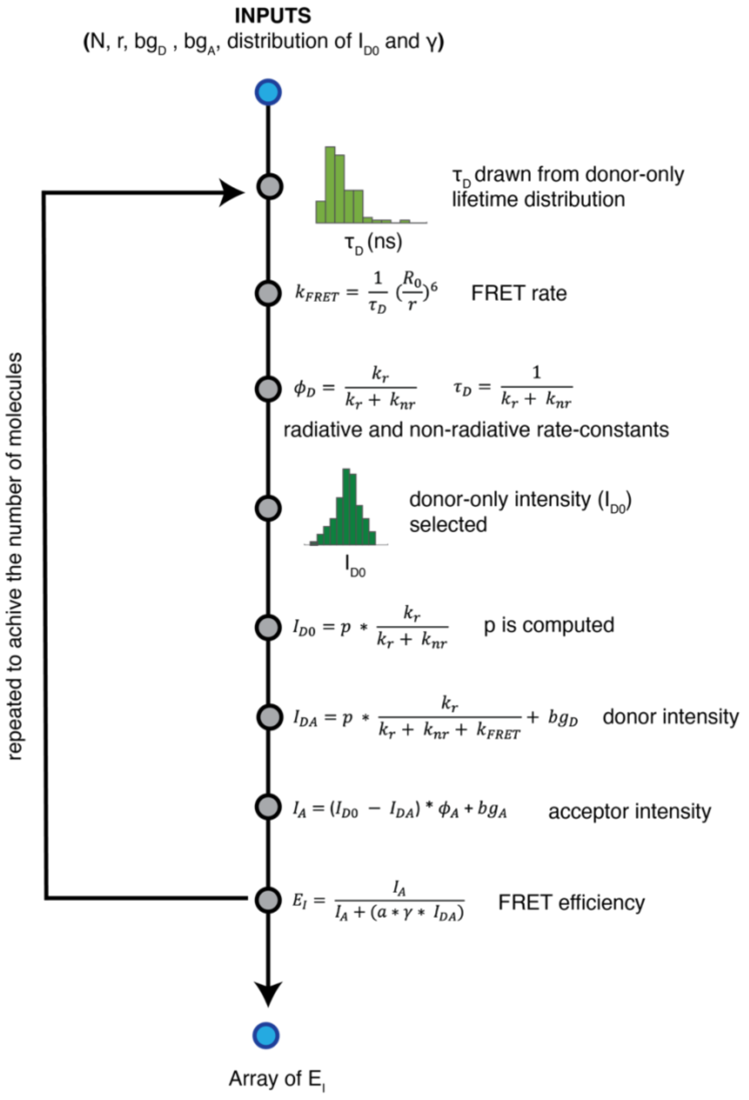


N=number of molecules to simulate, r= donor-acceptor distance, bgD,bgA=background in the donor and acceptor channels[3]. $\gamma$ is the correction factor.

In a perfectly aligned set-up, we can assume that fluorescence intensity drops in the donor channel due to FRET, is proportional to the increase in fluorescence intensity in the acceptor channel. Therefore, the acceptor fluorescence intensity ($I_A$) is given as:

$$I_A = (I_{D0} - I_{DA}) * \phi_A + bg_A \quad \text{(Eqn. 5)}$$

where, $\phi_A$=0.65, is the quantum yield for acceptor, $bg_A$ is the background in the acceptor channel.

Finally, the intensity-based FRET efficiency ($E_I$) is computed as:

$$E_I = \frac{I_A}{I_A + (a * \gamma * I_{DA})} \quad \text{(Eqn. 6)}$$

where $\gamma$ is the correction term arising due to different detection efficiency in the donor and acceptor channel and their quantum yields. In the simulation, we used a distribution of $\gamma$ for the same donor-acceptor pair obtained in a similar microscope set-up. $a$ is a parameter adjusted so that $E_I$ matches to the experimentally observed FRET efficiency for the mid-FRET construct and kept the same for the other constructs.

**Donor decay based FRET**

Intensity-based smFRET measurement is widely popular due to its straightforward implementation and analysis of the data. However, the fluorescence intensity of the donor and acceptor depends on many external factors, e.g., excitation power, microscope alignment, difference in detection efficiency in donor and acceptor channels, cross-talk etc. On the other hand, excited state lifetimes are not very sensitive to the factors mentioned above and therefore lifetime-based smFRET measurements are comparatively robust and accurate.

Once the donor molecule is excited with a $\delta$-pulse, the decay of excited state donor population ($[D]$) is governed by the equation below:

$$\frac{d[D]}{dt} = -(k_d + k_{FRET})[D] \quad \text{(Eqn. 7)}$$

Solution of the above equation gives,

$$[D] = D_0 e^{-(k_d + k_{FRET})t} = D_0 e^{-t/\tau_{DA}} \quad \text{(Eqn. 8)}$$

Where, $D_0$ is the initial population of the donor excited state after the $\delta$-pulse excitation. However, in practice, due to the finite width of the excitation pulse and the factors related to photon detections, optics and electronics, this decay is convolved with the instrument response function (IRF) of the system. Therefore, the donor decay is given as a convolution of IRF:

$$[D] = D_0 e^{-t/\tau_{DA}} \otimes IRF \quad \text{(Eqn. 9)}$$

In donor decay based FRET measurement approach, the fluorescence decay from the donor is fit to Eqn. 9 to extract $\tau_{DA}$. Due to low number of photons collected in the smFRET measurements, fits based on maximum likelihood estimation (MLE) give more robust results compared to the traditional chi-squared based fitting method. Therefore, $\tau_{DA}$ are obtained from the MLE fits. Once $\tau_{DA}$ is known, the FRET efficiency ($E$) can be computed from the familiar equation:

$$E = 1 - \frac{\tau_{DA}}{\tau_D} \quad \text{(Eqn. 10)}$$

**Supplementary Figure 14: Algorithm to simulate donor lifetime decay-based smFRET efficiency**

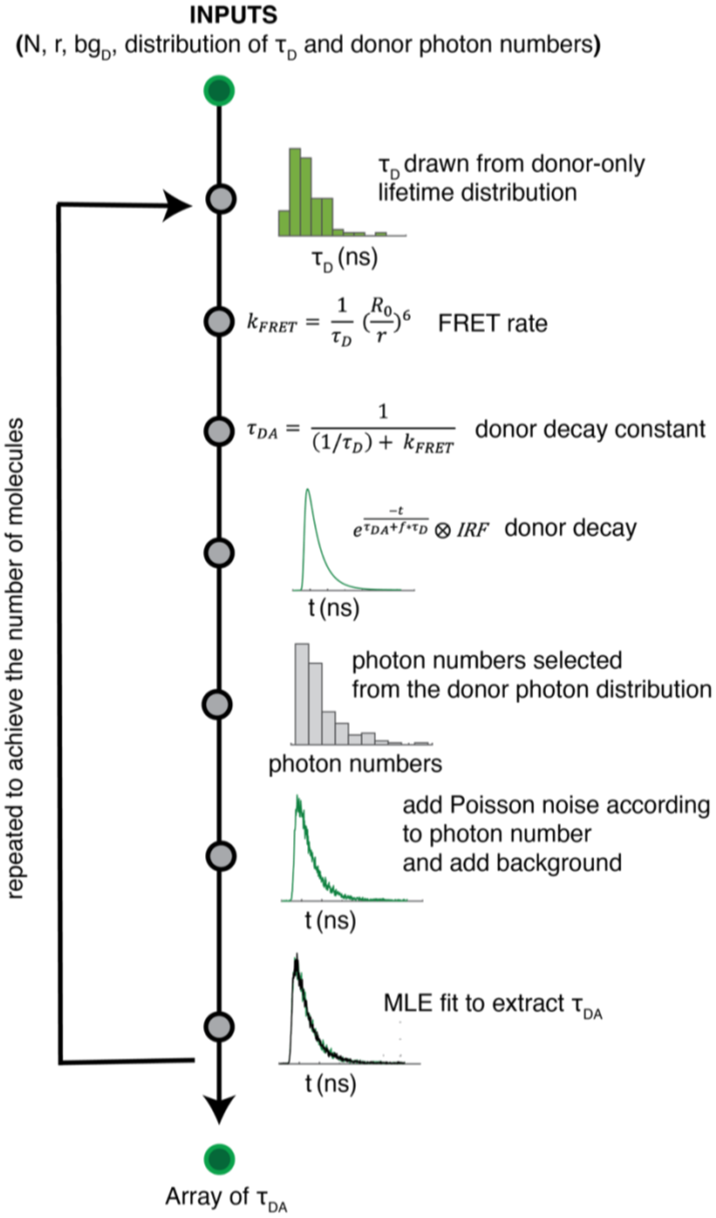


N=number of molecules to simulate, r=donor-acceptor distance, bgD=background in the donor channel.

Figure 12 demonstrate the algorithm to generate smFRET efficiency from the donor decay approach. Please note that to obtain the donor decay, the exponential part of the Eqn. 9 has been modified to:

$$e^{-\frac{t}{\tau_{DA}+f*\tau_D}} \quad \text{(Eqn. 11)}$$

Additional $f * \tau_D$ term in the exponent is added to account for the longer lifetime decay observed in the experimental data, particularly when the signal-to-noise ratio is low. We assume that this longer decay is due to the donor-only decay spill-over arising from either multiple donor in the construct or excitation of nearby donor molecules. We modeled $f$ as:

$$f = a * \frac{bg_D}{I_D + bg_D} \quad \text{(Eqn. 12)}$$

Where $bg_D$ and $I_D$ are the background and the fluorescence signal in the donor channel. $a$ is a constant set to match the mid-level FRET efficiency but kept the same value across the constructs. The Poisson noise added on the donor decay so that the area under the decay curve is equal to the photon number drawn from the total photon distribution in the donor channel. Finally, the donor decay is fit using MLE to extract $\tau_{DA}$ and $E$ is calculated from Eqn. 10.

**Supplementary Figure 15: Distribution of the donor decay lifetimes of the experiments and the simulations**

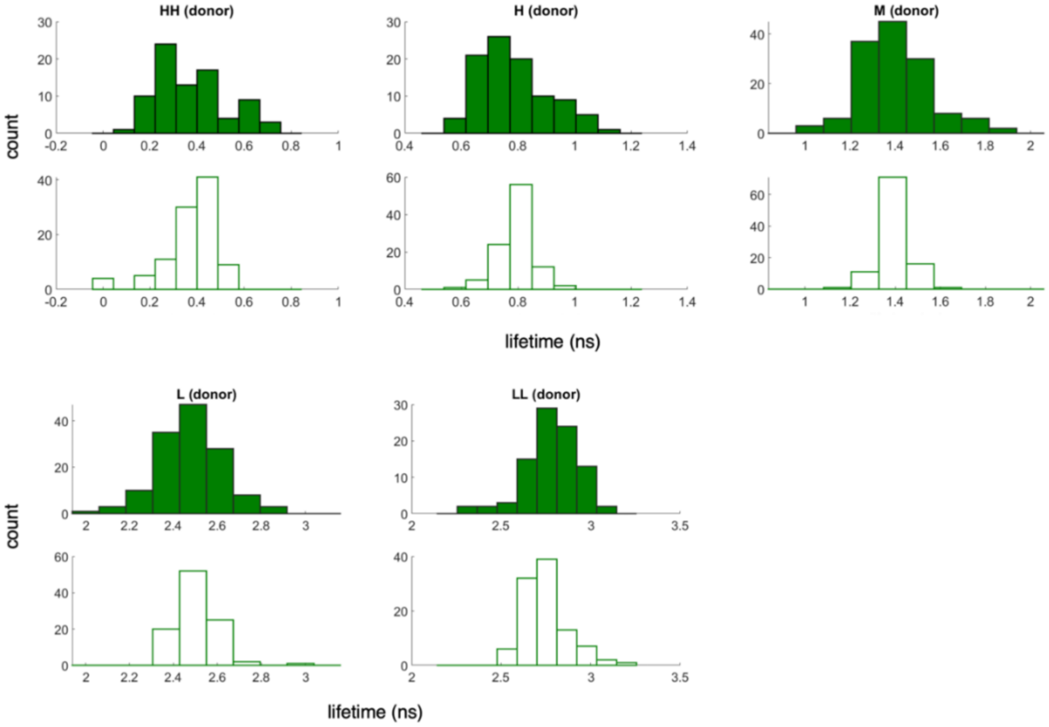


Distribution of the donor decay lifetimes from the experiments (solid histograms) and the simulations (hollow histograms) at different donor-acceptor distances.

Figure 13 shows the distribution of $\tau_{DA}$ obtained from experiments and as well as the simulations using donor decay method.

##### Acceptor rise based FRET

The decay of the acceptor is governed by the equation below:

$$\frac{d[A]}{dt} = -k_a[A] + k_{FRET}[D]$$
$$= -k_a[A] + D_0 e^{-t/\tau_D} \quad \text{(Eqn. 13)}$$

Solution of the above equation is given as[4].

$$[A] = \frac{D_0 k_{FRET}}{k_d + k_{FRET} - k_a} e^{-k_a t} - \frac{D_0 k_{FRET}}{k_d + k_{FRET} - k_a} e^{-(k_d + k_{FRET})t}$$
$$= \frac{D_0 k_{FRET}}{k_d + k_{FRET} - k_a} e^{-t/\tau_A} - \frac{D_0}{k_d + k_{FRET} - k_a} e^{-t/\tau_{DA}} \quad \text{(Eqn. 14)}$$

Please note that unlike the donor decay, the acceptor decay has both decay ($e^{-t/\tau_A}$) and a rise component ($-e^{t/\tau_{DA}}$). As the rise component has the term $\tau_{DA}$, fit of the acceptor rise can reveal the FRET efficiency (Eqn. 10).

**Supplementary Figure 16: Algorithm to simulate acceptor rise time-based smFRET efficiency**

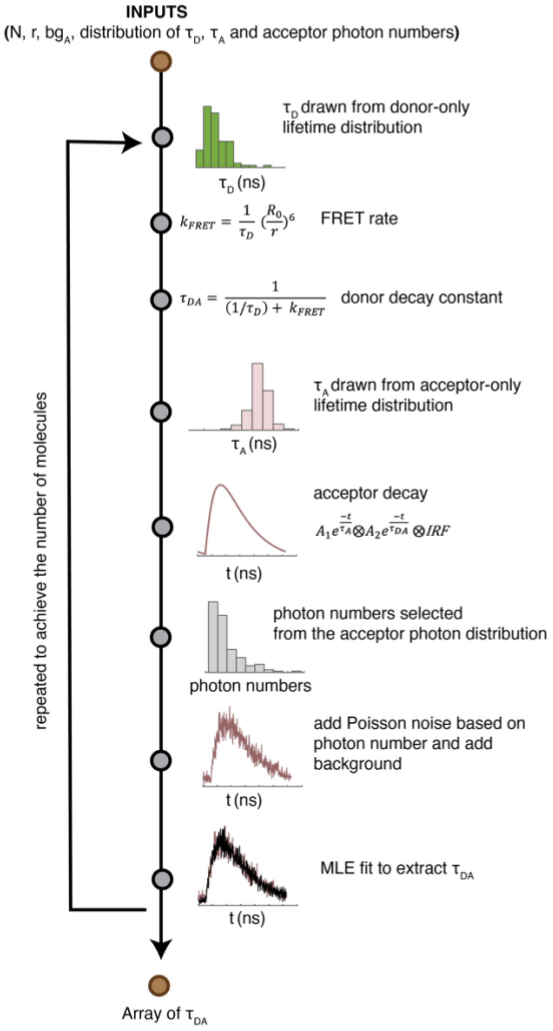


N = number of molecules to simulate, r= donor-acceptor distance, bgA = background in the acceptor channel.

Figure 14 demonstrate the algorithm to simulate the smFRET efficiency based on acceptor rise term. Although according to Eqn. 14, the amplitude of the decay and rise terms are equal, in practice, they can be different due to cross-talk and direct excitation of the acceptors[5]. Therefore, we model the acceptor decay using different amplitudes ($A_1, A_2$) of the decay and rise-terms. In the low FRET regime, the value of $\tau_{DA}$ approaches to $\tau_D$. Again, the donor-only and acceptor only lifetimes are similar in magnitudes (~3.5 ns). Therefore, in the low FRET regime, the two terms in the acceptor decay become indistinguishable and tends to cancel each other. Consequently, the Eqn. 14 become inaccurate to represent the acceptor decay as it approaches to zero. To avoid this situation, we modeled the acceptor decay as a convolution of $A_1 e^{-\frac{t}{\tau_A}}$ , $A_2\, e^{-\frac{t}{\tau_{DA}}}$ and IRF. Similar to donor decay, Poisson noise is added to the acceptor decay model so that the area under the decay curve becomes equal to the number of total photons selected from the total acceptor intensity distribution. Finally, the acceptor decay is fit using MLE method to extract $\tau_{DA}$ and FRET efficiency.

**Supplementary Figure 17: Distribution of the acceptor rise time of the experiments and the simulations**

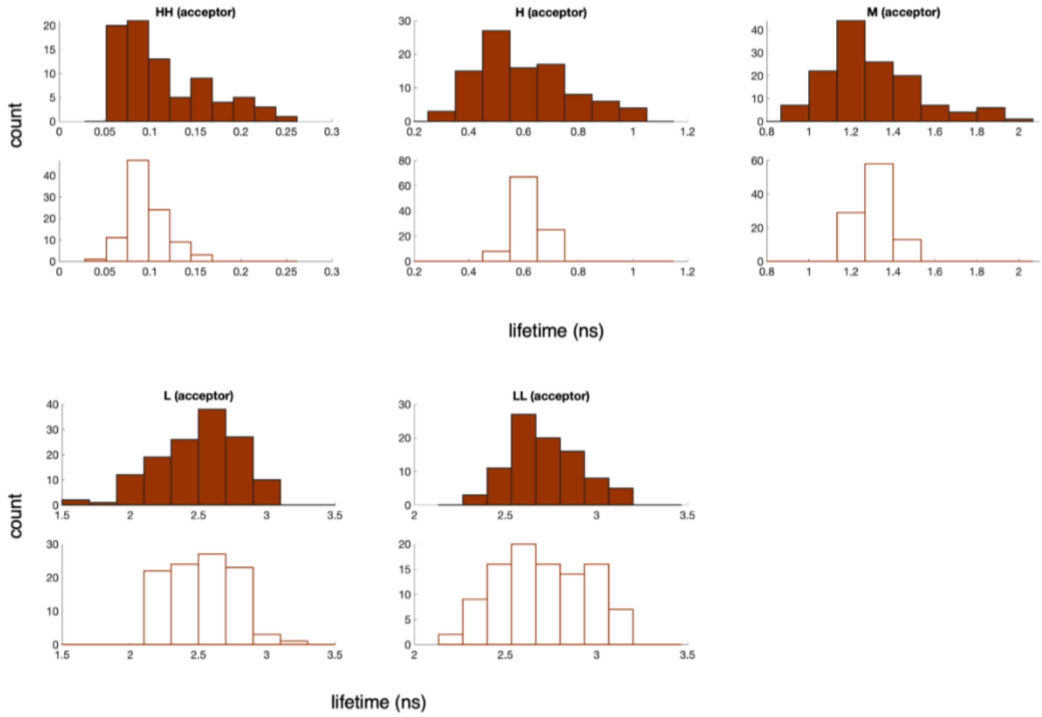


Distribution of acceptor rise time from the experiments (solid histograms) and the simulations (hollow histograms) at different donor-acceptor distances.

**Supplementary Table 8: Summary of the FRET efficiencies of experiments and the simulations using donor decay, acceptor rise and intensity-based methods.**

| Value<br>Error<br>(bootstrap) | high-high FRET | high FRET | mid FRET | low FRET | low-low FRET |
|---|---|---|---|---|---|
| smART-$E_{FRET}$ | 0.97<br>0.000 | 0.81<br>0.003 | 0.59<br>0.004 | 0.21<br>0.012 | 0.16<br>0.013 |
| smART-Distance (nm) | 3.58<br>0.000 | 5.05<br>0.000 | 6.01<br>0.010 | 7.97<br>0.080 | 8.46<br>0.120 |
| Donor decay-$E_{FRET}$ | 0.88<br>0.007 | 0.76<br>0.003 | 0.57<br>0.002 | 0.22<br>0.002 | 0.14<br>0.002 |
| Donor decay-Distance (nm) | 4.57<br>0.010 | 5.30<br>0.000 | 6.11<br>0.000 | 7.92<br>0.010 | 8.63<br>0.020 |
| Intensity-$E_{FRET}$ | 0.90<br>0.011 | 0.80<br>0.012 | 0.61<br>0.018 | 0.21<br>0.016 | 0.10<br>0.009 |
| Intensity-Distance (nm) | 4.42<br>0.010 | 5.08<br>0.010 | 5.95<br>0.030 | 7.95<br>0.100 | 9.21<br>0.140 |

The numbers in the parenthesis are the standard deviations. The number of molecules used in the simulations is ~100 which is comparable to the experiments.

#### Part 3: Molecular dynamics simulations of the SSB-DNA complex

To facilitate the interpretation of smART FRET experiments, we carried out atomistic molecular dynamics simulations of the SSB-DNA complex. A structural model of the complex was built with homology modeling based on two experimentally solved E. coli SSB crystal structures: 1EYG and 1QVC by Maffeo et al.[6] We solvated the protein-DNA complex structure in a dodecahedron box with a total of 25185 TIP3P water molecules. The box size was set to ensure a minimum distance of one nm between the solute and the boundary. Sodium and chloride ions were added to neutralize the system and reproduce the three salt concentrations studied experimentally: 100mM, 150mM, and 400mM. We carried out an independent simulation for each of the three systems at different salt concentrations under constant pressure and temperature (NPT) using the Parrinello-Rahman barostat[7] and the velocity-rescaling thermostat[8]. The Particle Mesh Ewald method[9] was used to evaluate electrostatic interactions, with a grid spacing of 0.12 nm. All simulations lasted 2.2 $\mu s$ or longer with a timestep of 2 fs. The first 1.2 $\mu s$ of each trajectory was discarded as equilibration, and configurations collected from the remaining segments at every one ns were used for structural analysis. When computing the distance between nucleotides 1 and 70 to compare with experimental values, the distance between the geometric centers was used.

**Supplementary Figure 18: Traces of SSB-DNA molecular dynamics.**

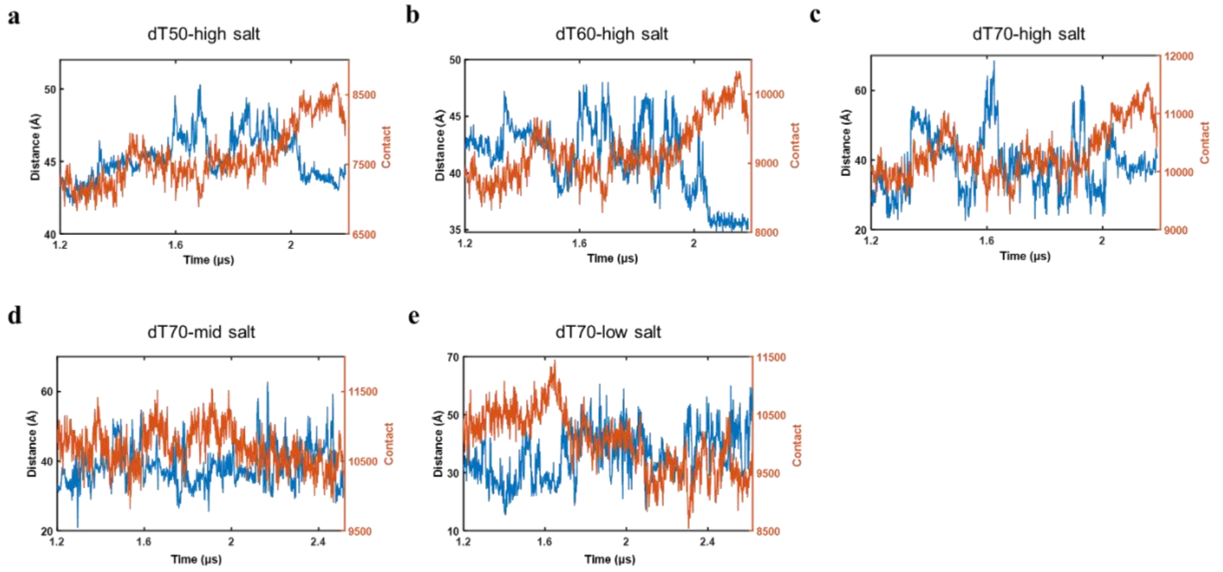

**Supplementary Figure 19: Bi-modal fitting of SSB-DNA (dT70) molecular dynamic simulation in 400mM NaCl.**

The set of simulated distances was binned into a histogram (bin = 31) and fitted to a bi-modal distribution.

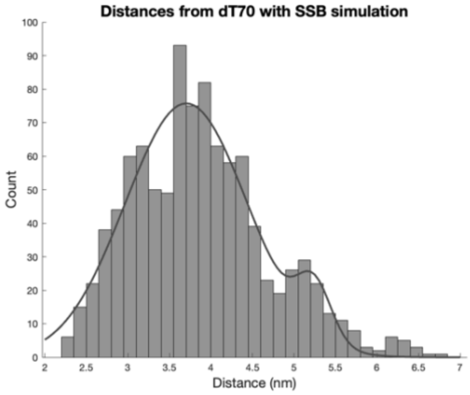


**Supplementary Table 9: Summary of SSB-DNA molecular dynamic simulation results.**

| Value<br>Error<br>(bootstrap) | high salt<br>dT50 | high salt<br>dT60 | high salt<br>dT70 | mid salt<br>dT70 | low salt<br>dT70 |
|---|---|---|---|---|---|
| Distance (nm) | 4.53<br>0.009 | 4.13<br>0.020 | 3.88<br>0.056 | 3.85<br>0.029 | 3.60<br>0.044 |

**Supplementary Table 10: Summary of bi-modal fitting of SSB-DNA (dT70) molecular dynamic simulation in 400mM NaCl.**

The distances and errors were extracted from the mean and sigma of the bi-modal fitting. The ratio of tight and loose state is calculated from the amplitude of the tight to open state.

| Value<br>Error | Tight<br>conformation | Loose<br>conformation |
|---|---|---|
| Distance (nm) | 3.73<br>0.74 | 5.24<br>0.21 |
| Amplitude | 73.4 | 16.3 |
